# Individualized non-invasive brain stimulation engages the subgenual anterior cingulate and amygdala

**DOI:** 10.1101/503441

**Authors:** Desmond J. Oathes, Jared Zimmerman, Romain Duprat, Seda Cavdaroglu, Morgan Scully, Benjamin Rosenberg, Matthew W. Flounders, Hannah Long, Mark Elliott, Gavriella Shandler, Russell T. Shinohara, Kristin A. Linn

## Abstract

Brain stimulation is used clinically to treat a variety of neurological and psychiatric conditions. The mechanisms of the clinical effects of these brain-based therapies are presumably dependent on their effects on brain networks. It has been hypothesized that using individualized brain network maps is an optimal strategy for defining network boundaries and topologies. Traditional non-invasive imaging can determine correlations between structural or functional time series. However, they cannot easily establish hierarchies in communication flow as done in non-human animals using invasive methods. In the present study, we interleaved functional MRI recordings with non-invasive transcranial magnetic stimulation in the attempt to map causal communication between the prefrontal cortex and two subcortical structures thought to contribute to affective dysregulation: the subgenual anterior cingulate cortex (sgACC) and the amygdala. In both cases, we found evidence that these brain areas were engaged when TMS was applied to prefrontal sites determined from each participant’s previous fMRI scan. Specifically, after transforming individual participant images to within-scan quantiles of evoked TMS response, we modeled the average quantile response within a given region across stimulation sites and individuals to demonstrate that the targets were differentially influenced by TMS. Furthermore, we found that the sgACC distributed brain network, estimated in a separate cohort, was engaged in response to sgACC focused TMS and was partially separable from the proximal default mode network response. The amygdala, but not its distributed network, responded to TMS. Our findings indicate that individual targeting and brain response measurements usefully capture causal circuit mapping to the sgACC and amygdala in humans, setting the stage for approaches to noninvasively modulate subcortical nodes of distributed brain networks in clinical interventions and mechanistic human neuroscience studies.

## Introduction

Resting functional MRI seed-based connectivity is a standard approach for summarizing brain ‘networks’ that are putatively functionally linked and thought to subserve complex and basic mental operations. Our recent work suggests that evoking activity with transcranial magnetic stimulation (TMS) in a surface accessible node of intrinsic networks largely recapitulates within and between-network correlation maps (1). Emerging approaches in fMRI connectivity promote making greater use of individual topographic maps before aggregating group data so as to capture more accurately functional landscapes from each individual that are spatially variable across subjects but reproducible within subjects (2, 3). In the present study, we approached ‘individualization’ in brain stimulation in both target determination and measurement of brain responses to stimulation. Targets were individualized using each individual’s resting fMRI scan by seeding a deep subcortical region of interest and mapping prefontal cortical areas of highest timeseries correlation with the target for that individual, i.e., ‘resting fMRI guided TMS’.

Previous work used individual quantile ranks to describe canonical regional bins in cortical thickness and resting fMRI signal amplitude to demonstrate age-related topological patterns (4). We explored quantile ranks calculated within individual interleaved TMS/fMRI scans across voxels, which has not been previously reported to our knowledge. The motivation for using quantiles was based on interest in the relative distribution of TMS evoked brain activation for an individual brain as a basis for effective targeting of brain networks and systems. This strategy has direct clinical application as a basis for personalized medicine.

The current investigation focused on stimulation of accessible regions of prefrontal cortex hypothesized to influence deep brain structures related to affective disturbance: the amygdala and the subgenual anterior cingulate cortex (sgACC). The amygdala is essential to a range of emotional processes and is known to be dysregulated in affective disorders such as posttraumatic stress disorder, social phobia and depression (5, 6). A large body of neuroimaging work has identified prefrontal regions thought to potentially regulate amygdala activity (7, 8) and also reliably shown to be dysfunctional in clinical populations (9). In a variety of non-human primate studies, amygdala functional connectivity studies in humans, and a recent diffusion tractography study, multiple subregions of prefrontal cortex stand out as potential targets by which TMS might causally influence the amygdala (10-12). The importance of the sgACC is highlighted by many neuroimaging studies of clinical depression as well as negative mood induction (13-16). Based on evidence that optimal responses from deep brain stimulation implants for treating depression seem to occur with electrode placement proximal to white matter pathways innervating the subgenual cingulate (17-19), ‘connectivity’ to this target is a priority in defining possible stimulation targets for non-invasive brain stimulation. In the clinical domain, when repetitive TMS treatment is delivered to regions of prefrontal cortex with higher magnitude resting time series correlation with sgACC, the treatment tends to work more effectively (20, 21). However, the neuroimaging, clinical TMS and animal studies do not clearly prove that there are prefrontal cortical locations that causally influence the amygdala or sgACC in humans. In one recent report of dorsolateral prefrontal cortex stimulation with online fMRI, there was on average no sgACC response (22) though one subject in particular may have activated the sgACC.

To test for the possibility that a targeted approach might causally, on average, engage the sgACC and amygdala, we interleaved TMS single pulses with fMRI recordings which, given the slow rise time of the BOLD signal response to an ‘event’ (TMS here), allows a brain-wide causal activation map to be defined for any surface brain stimulation target. This is achieved by probing the accessible cortical site and measuring fMRI BOLD signal downstream in response to stimulation. In contrast to our previous work using atlas based targeting (1), here we made individual brain targets based on each participant’s initial resting fMRI scan that were used to generate TMS stimulation sites for the subsequent TMS/fMRI scan. We believe this is an important step forward that capitalizes on recent work suggesting that network representations in the brain are highly individual, as well as evidence that different networks may exist at the same anatomical location across subjects (23).

There is justification for looking at the sgACC and amygdala regions individually, though both of these regions communicate with distributed networks likely relevant to complex mental operations subserving emotion and its dysregulation in affective illness. Therefore, we generated network masks for the sgACC and amygdala individually using our processing pipeline applied to a large independent healthy cohort from a publicly available source.

Among sites accessible to TMS while participants laid on their backs in the scanner, we considered areas of left hemisphere prefrontal cortex with high resting connectivity to the subcortical target of interest as stimulation sites. There were multiple sites for each participant for each target (each at least 1.47cm apart and none within 2cm of another target immediately preceding or following it in sequence). For each participant, across 1-2 TMS/fMRI sessions, we stimulated multiple accessible targets for the sgACC and for the amygdala (see Figure 1). By including multiple stimulation sites for each subcortical target, we sought to determine whether resting fMRI targeted TMS is effective, generally, rather than confined to a particular sub-region of prefrontal cortex. Our priority was to establish the degree to which the downstream target was engaged, since this was the seed for the connectivity maps used to choose the stimulation sites. Similarly, we aimed to establish the degree to which distributed networks derived from those seed maps were engaged with TMS. In follow-up, exploratory analyses, we investigated sensitivity to statistical modeling choices, specificity of our regions of interest, the degree of sgACC network response excluding overlapping DMN regions, whole brain responses by target, and arousal contributions to the evoked responses.

**Figure 1.**
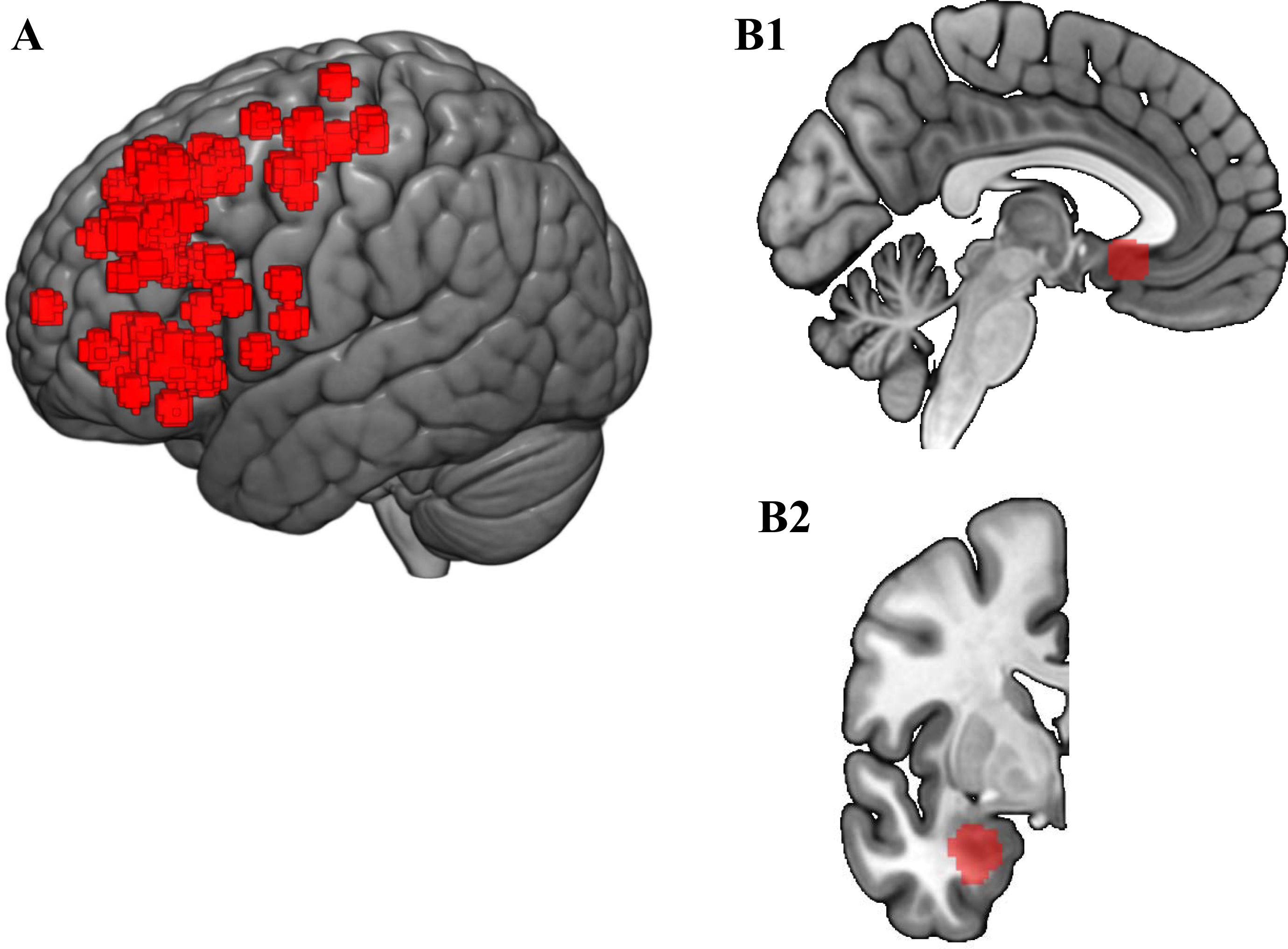
Sites of stimulation and downstream targets. (**A**) All stimulation sites. (**B**) Subcortical regions seeded to generate functional MRI individual correlated surface accessible brain regions for stimulation in A. **B1** is the subgenual anterior cingulate cortex at MNI x= −4; **B2** is the left basolateral amygdala at MNI y= −2.

## Materials and methods

### Participants

All participants gave consent for the experiment according to the Declaration of Helsinki and the study was approved by the University of Pennsylvania IRB. Participants ranged from age 22-42 (mean 28.71, standard deviation 4.95); had Bachelor’s to Doctoral education; had no history of a neurological or psychiatric condition; and were not taking any psychoactive medications. There were 8 male and 6 female participants. In total, 71 TMS/fMRI datasets (sites) were acquired across 1-2 sessions per individual.

### Equipment

MRI data were acquired on a 3 Tesla Siemens Prisma scanner using one head coil for baseline resting and structural MRI (32 channel; Erlangen, Germany) and another for TMS/fMRI (RAPID quad T/R single channel; Rimpar, Germany) in order to accommodate the custom built TMS coil holder and TMS coil. TMS was delivered using an MRI compatible Magventure MRI-B91 air cooled TMS coil connected to a Magpro X100 stimulator (Magventure; Farum, Denmark). Neuronavigation through a stereotaxic system (Brainsight; Rogue Research, Montreal, Quebec, Canada) matching fiduciary points from the participant with MRI images through a Polaris optical position Vicra camera allowed marking the stimulation sites on a lycra swim cap. The location for each site was based on individual functional connectivity values over which the TMS coil was placed in the MRI. A dedicated windows PC installed with E-prime 2.0 (Psychology Software Tools, Sharpsburg Pennsylvania USA) was used to gate TMS pulse delivery as well as MRI scans between pulses that each were triggered via TTL pulses through a parallel port with unique pin assignments for each device.

### Baseline MRI acquisition

For each subject, a resting state fMRI scan with phase encoding direction anterior to posterior was acquired (TR=720ms, TE=37ms, FA=52°, FOV=208mm, 2×2×2mm voxels, 72 interleaved axial slices with no gap, 600 measurements). For that sequence, subjects were instructed to keep their eyes open and remain as still as possible. Structural data were also acquired and consisted of a high-resolution multi-echo T1-weighted MPR image (TR=2400ms, TI=1060ms, TE= 2.24ms, FA=8°, 0.8×0.8×0.8mm voxels, FOV= 256mm, PAT mode GRAPPA, 208 slices).

### TMS/fMRI data

Interleaved TMS/fMRI images were acquired with a TR of 2000ms and a TE of 30ms (FA=75°, FOV=192mm, 3×3×4mm voxels, 32 interleaved axial slices, 174 measurements). Each volume was gated by TTL triggers from Eprime and with a 400ms gap between volumes to allow interleaved TMS pulses to be delivered halfway through the gap (at 200ms). This avoids known T1 contamination by simultaneous RF and TMS pulses (24) but given slow BOLD rise time, effectively captures TMS evoked brain responses. Single pulses were interleaved in 12 mini-blocks of 7 stimulations separated by 1 TR and including 0, 1, or 2 ‘catch trials’ (random order and block) during which spacing was separated by an extra TR with no TMS pulse to avoid easy prediction of TMS delivery by participants. The mini-blocks were themselves separated by 7 TRs and 71 total stimulations were given per site over 174 volume acquisitions.

### MRI processing

For processing resting fMRI data, the first 10 volumes were discarded for T1 equilibrium, and then an automated removal of motion artifact designed to maintain fMRI autocorrelation structure using ICA-AROMA (25) was applied. Nuisance regression, residualizing for white matter and CSF signal, was implemented in FSL 5.08 (FMRIB Oxford, UK), followed by bandpass filtering 0.008-0.1 Hz and 6mm kernel FWHM smoothing. Boundary based registration following FSL FAST tissue segmentation used 6 degrees of freedom to coregister T1 to functional scans and FSL FNIRT default settings were used for nonlinear warps of fMRI data to the 2mm MNI152 average template (26). The inverse of this process moved seed regions to native fMRI space in one step, and functional connectivity for each seed was calculated in native space, resulting in Pearson correlation maps that were transformed to z-scores using the Fisher rto-z equation. Inverting the T1 to functional transform placed the fMRI targets in native T1 space for neuronavigation that were also visually verified. TMS/fMRI data analysis removed two initial volumes for T1 equilibrium and then regressed TMS events convolved with a hemodynamic response function (SPM12 double gamma) on the fMRI time series with regressors of no interest and six motion parameters derived from a 6 degree of freedom linear (FSL FLIRT) transform that coregistered fMRI images to the middle volume acquired. The resulting contrast estimates were subjected to additional analysis and statistical procedures described below.

### TMS targets

For both sgACC and amygdala targets, resting FC z-score values of absolute value 0.25 or greater were considered for targeting. For some sites, the FC values for both targets exceeded this threshold and so were included in both sgACC and amygdala focused analyses. For sgACC targets, there were 41 sites (TMS/fMRI runs), including 24 that did not have supra threshold amygdala connectivity. For the amygdala (BLA), there were 44 sites, of which 26 did not have supra threshold sgACC connectivity. For sgACC targets, 7 had negative connectivity FC values with the sgACC and for amygdala targets, 6 had negative FC values with the amygdala. For both the sgACC and BLA, all negatively correlated sites had a within-network TMS evoked quantile response that was within 1 SD of the average within-network quantile response of the positively correlated sites. For the ROI analyses, all negatively correlated sites had a within-BLA TMS evoked quantile response within 1 SD of the average within-BLA quantile response of the positively correlated sites, with the exception of one value that was within 1.5 SDs. Similarly, for the sgACC target, one value was within 1.6 SDs, another within 1.3 SDs, and the rest were within 1 SD of the positively correlated site average.

The TMS coil was positioned with the coil handle facing backwards for a posterior to anterior induced current. The FDI (first dorsal interosseuous) or APB (abductor pollicis brevis) of participant’s dominant/right hand was used (whichever most clearly responded) as target muscles for determining motor threshold based on visual observation (5/10 trials with a noticeable resting muscle twitch). Stimulation intensity was then set to 120% motor threshold for all stimulation delivered to that participant.

The sgACC seed was based on an average MDD associated abnormality across fourteen neuroimaging studies and shown to change in connectivity following rTMS treatment (27) shifted just to the left of midline ipsilateral to the stimulation sites and also anatomically well centered to Brodmann 25 at MNI −2, 18, −8 (Figure 1B1). As in previous amygdala subregional fMRI studies (28), the basolateral amygdala (Figure 1B2) was defined using a probability map with threshold 40% for the BLA from a common histological atlas (29), and voxels were retained only if they exceeded the probability threshold for adjoining subregions (centromedial amygdala, CMA; superficial amygdala, SF) (30).

The gray matter mask was based on visual confirmation using FSL version 5.0.8 with an arbitrary value of ‘100’ and up based on visual inspection on the 152 average T1 gray matter tissue prior for a mask in standard space.

### Network mask creation

One hundred and twenty seven healthy adult participants with low head motion (<2.0 mm) from a prior release of the NKI ‘extended’ resting fMRI and structural T1 database were used to define normative atlases for each of the sgACC (Figure 2) and amygdala (Figure 3) seed regions, as in our prior work (31). The full sample z-scores thresholded respectively above 0.3 and 0.2 generated voxelwise maps that were given a count of ‘1’ for every subject who had an absolute value z-score that exceeded this threshold. If the count for that voxel exceeded 96 (out of a possible 127 subjects; >75%), the voxel was retained for the next step. The values for the remaining voxels were used to calculate a median image for the whole sample and a standard deviation, both of which were used to generate an effect size (median/standard deviation) required to exceed 6.0 and with a minimum cluster size of 10 voxels (2 mm^3^) to be retained in the respective network mask. All other processing steps were identical to those from data obtained locally.

**Figure 2.**
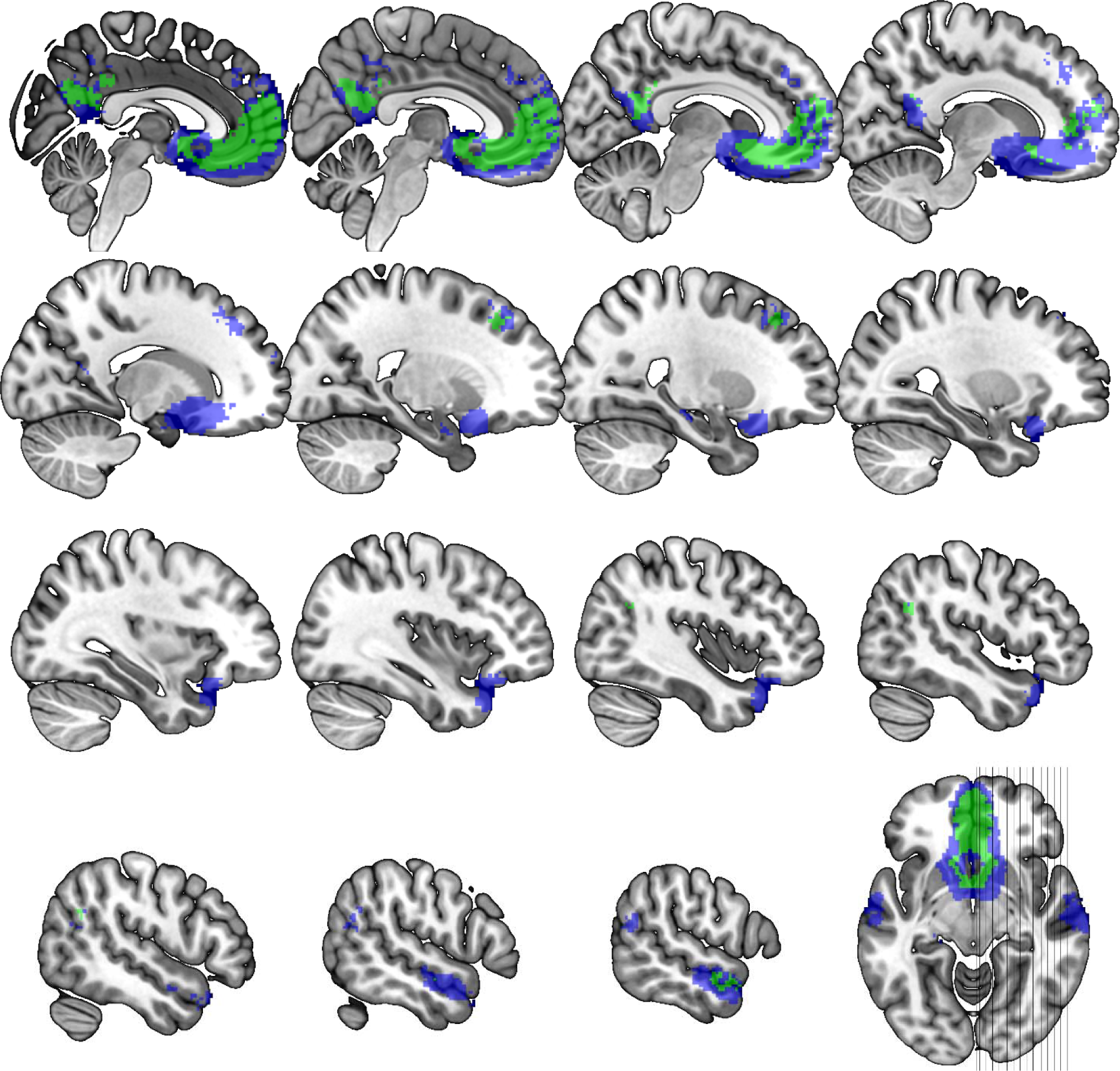
Subgenual cingulate network. Using an independent data set, the network mask shown was created seeding the primary (green) or FreeSurfer (Blue) defined sgACC and calculating functional connectivity (correlation) with z≧0.3 in 96+/127 subjects, effect size >6.0 and cluster ≧ 2mm3 in 2mm MNI standard space (excluding seed). Slice wise views are represented starting at MNI x=2 and moving out in 2mm steps with an extra step between rows until the final x=58 image.

**Figure 3.**
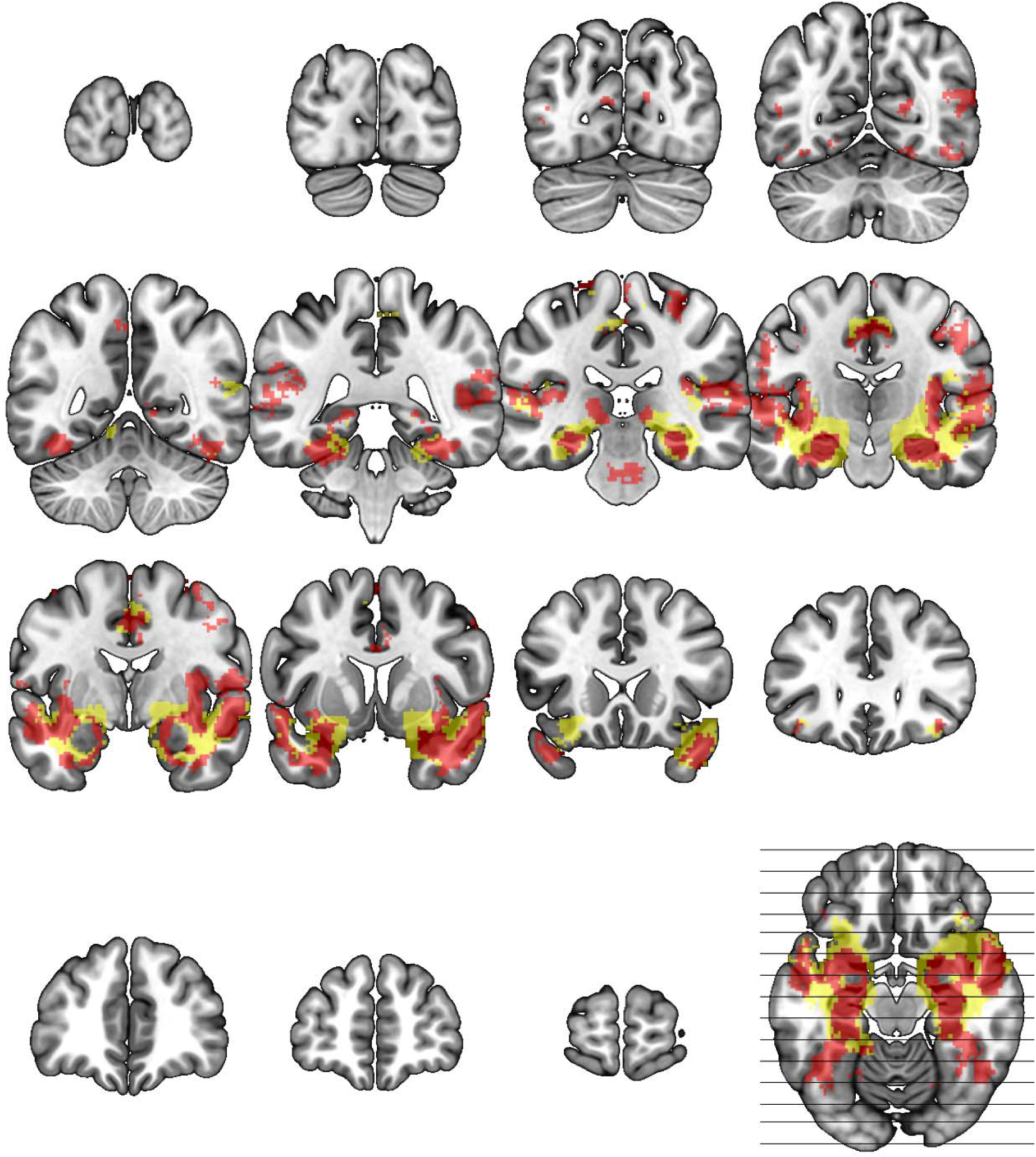
Basolateral and full amygdala network. Using an independent data set, the network mask shown was created seeding the BLA (red) or FreeSurfer amygdala (yellow) defined regions of interest and calculating functional connectivity (correlation) with z≧0.3 in 96+/127 subjects, effect size >6.0 and cluster ≧ 2mm3 in 2mm MNI standard space (excluding seed). Slice wise views are represented starting at MNI y=-98 and moving forward in 12mm steps until the final y=66 image.

### FreeSurfer ROIs

Standard space masks for the amygdala and sgACC were used from FreeSurfer (32). The MNI152 1mm template brain was run through FreeSurfer’s ‘recon-all’ pipeline to produce subcortical segmentations and cortical parcellations in MNI space. The amygdala ROI is from FreeSurfer’s implementation of the Harvard-Oxford subcortical atlas (33), and the sgACC ROI is the subcallosal gyrus ROI from the Destrieux atlas (34). In addition to using these alternative regions to test the specificity of the TMS evoked results to the primary subcortical targets, these seeds were used to generate alternative functional connectivity network maps as shown in the overlays in Figures 2 and 3. The overlap in the primary and FreeSurfer defined regions/seeds are also shown in Supplementary Figure 1.

### Statistical methods

Contrast estimates from the TMS evoked fMRI data demonstrated non-comparable levels of variability across participants (Supplementary Figure 2), which motivated our transformation of the contrasts to within-scan quantiles before performing group-level tests. From first level GLM contrast estimates in template space, the gray matter from each individual image (TMS stimulation site) was converted into an image of quantiles by ranking gray matter voxels from lowest to highest intensity and dividing by the total number of gray matter voxels. The quantile technique effectively normalized each individual’s brain response to TMS to a common domain, similar to allowing for subject-specific random effects in a mixed model. As a result, spatially distributed patterns of evoked TMS response could be pooled across stimulation sites and subjects without being obscured by large differences in global responses to TMS throughout the brain.

In our primary analysis, we tested whether the spatial patterns observed in the TMS activation quantile images had any correspondence across subjects and stimulation sites. For specific regions and networks of interest, we computed the average within-region quantile for each participant and tested whether the across-subject, across-site average of these summaries was different from the median of 0.5, which is the value that would be expected if quantiles across the brain had no coherent spatial pattern. Masks were used to extract the dependent variable, i.e., the average quantile within the region of interest minus 0.5 (i.e., we centered by the median). To account for possible correlation among repeated measures within subjects, we used generalized estimating equations (GEE), which are a semi-parametric extension of generalized linear models (GLMs) for non-independent data (35). In light of the number of subjects in our study (N=14), we opted to use jackknife standard error estimates (36), which have been shown to be less biased than the usual robust standard errors (37) in small samples (38, 39). All models controlled for pain ratings (z-scored within-subject across stimulation sites), centered age, and centered head motion (mean absolute value of root-mean-squared, 6 degrees of freedom calculated with fMRI image realignment). We tested whether the intercept in the GEE model was significantly (p<0.05) different from zero. Since we centered the dependent variable by 0.5 and centered the covariates in the model, a significant intercept can be interpreted as an average BOLD increase or decrease, depending on the sign, relative to median-level activation in the brain for a subject with average values of the covariates. For all primary network and region analyses, we used the identity link function and specified an exchangeable working correlation structure.

We confirmed the interpretability of the quantile responses by validating above- and below-median average quantile response with the direction of average TMS evoked responses in the 1^st^ level contrast estimates in fMRI BOLD. Given the level of correspondence of the measurements (demonstrated in a separate section below), we operationally label the negative and positive GEE intercept parameter estimates as ‘negative’ or ‘positive’ BOLD responses in the Results section. As a sensitivity analysis, we demonstrated robustness of the results by re-fitting all models with an independent correlation structure and qualitatively assessing changes in parameter estimates and significance levels. We also repeated the analyses additionally adjusting for heart rate and respiration measures, z-scored within subject, to quantify the effects of these covariates on the quantiles of evoked brain responses.

Finally, we performed an exploratory voxelwise analysis using the 44 amygdala target quantile images. We fit a GEE at every voxel (within the amygdala subregion mask) to account for repeated measures within subject. More specifically, the dependent variable at a given voxel was an indicator with value 1 if the quantile at that voxel was greater than the median quantile, 0.5, and 0 if it was less than the median. The collection of indicators at the voxel made up the response vector, which was regressed using the GEE on pain (z-scored within subject), centered age, and centered motion. We used the logit link function and specified an exchangeable working correlation structure. Since the assumptions for performing other family-wise error correction thresholding are not met when using a GEE, we corrected for multiple testing by controlling the false discovery rate (FDR) at q<0.05 within the amygdala submask. In the Supplement, we provide arbitrarily thresholded, unadjusted p-values from voxelwise GEE analyses of the entire gray matter separately for sgACC and amygdala target quantile for illustration purposes and to guide potential future projects.

### Data availability

The data supporting the study findings are available from the corresponding author upon reasonable request.

## Results

Participants had between four and fourteen TMS-fMRI scan runs across one or two scanning sessions. In each TMS-fMRI session the TMS coil was placed on the scalp at a location determined immediately prior to the scan in a TMS neuronavigation session to target prefrontal sites with strong functional connectivity to either the amygdala or sgACC. All models controlled for pain ratings (z-scored within-subject across stimulation sites), age, and head motion (mean absolute value of root-mean-squared, 6 degrees of freedom calculated with fMRI image realignment).

### Both amygdala and subgenual cingulate targeted TMS influenced their proposed target regions

We used the basolateral amygdala (BLA) as a seed to target prefrontal stimulation but anticipated that interconnected subregions of the amygdala might have also responded to TMS given their proximity and reciprocal functional connections. When testing the ipsilateral BLA, we did not establish evidence for activation, i.e., significantly greater than the median (0.5) quantile, with a liberal >=30% BLA probability mask from histological maps (29) averaged over the entire mask (see Figure 4; parameter estimate (PE)=0.003, Wald χ^2^=0.01, p=0.94). Exploring a smaller 50% BLA map did not change the findings (PE=0.007, Wald χ^2^=0.05, p=0.82). In the centromedial and superficial amygdala, we found significant activation in both subregions (CMA 30% PE=0.054, Wald χ^2^=4.25, p=0.039; SF 30% PE=0.055, Wald χ^2^=4.54, p=0.033) and pain was not significant in either model as a covariate. Using an independent amygdala ROI from the Harvard-Oxford subcortical atlas (33) yielded parameter estimates in the same direction but did not yield significant results (left amygdala PE=0.038, Wald χ^2^=1.08, p=0.300; full amygdala PE=0.014, Wald χ^2^<0.01, p=0.970), suggesting that specific subregions are particularly influenced by this amygdala targeted TMS approach.

**Figure 4.**
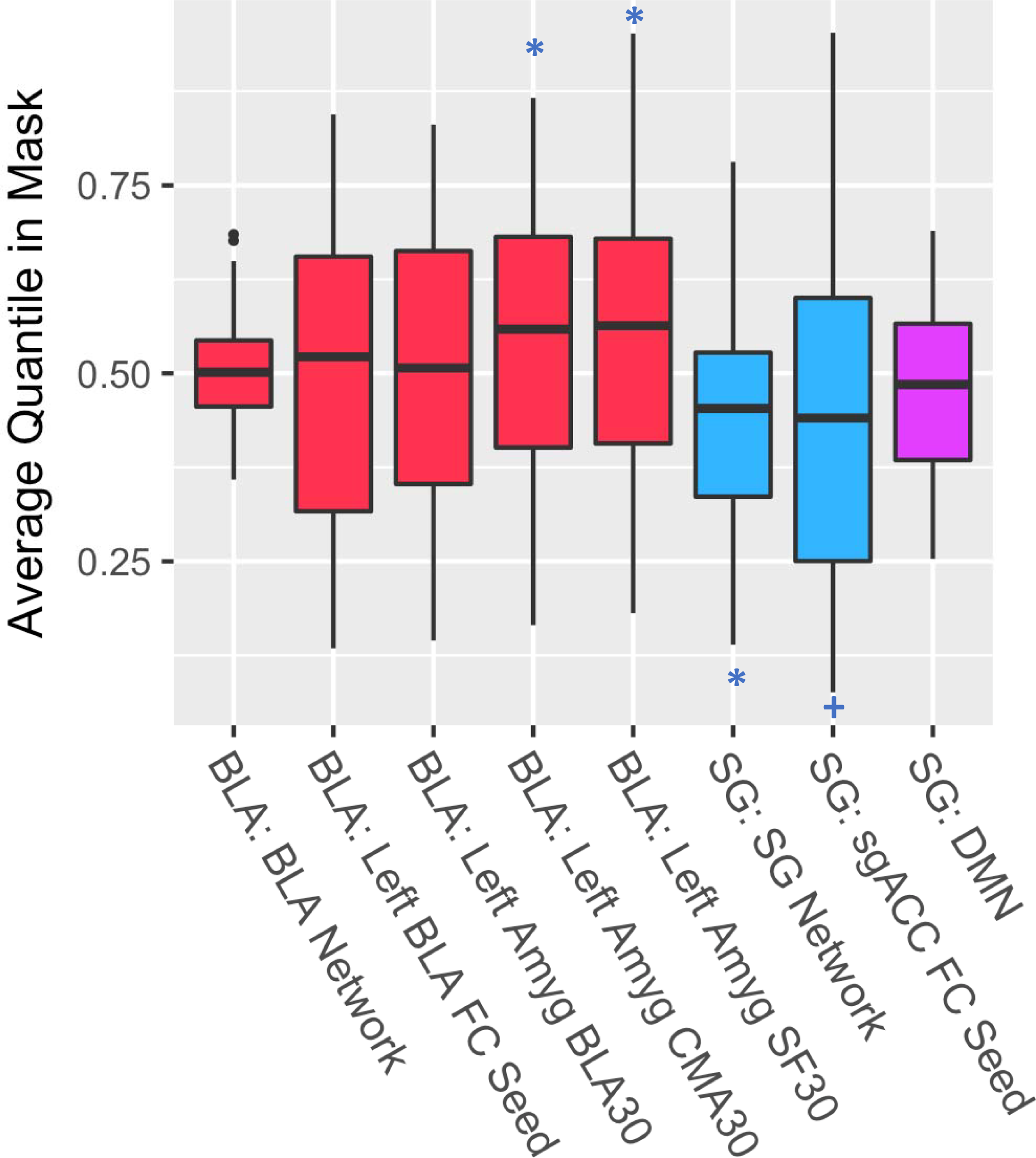
Primary results of generalized estimating equations controlling for age, TMS discomfort rating and head motion. Solid lines in the bars represent the median and the box hinges are the 25^th^ and 75^th^ percentiles. Whiskers are at the min/max but no further than 1.5* IQR (interquartile range). All results are centered at the 0.5 average brain response so quantiles above represent ‘lower’ and quantiles above represent ‘higher’ than the brain average TMS evoked response. ‘BLA’ and ‘SG’ labels before the colon represent the stimulation target of Basolateral Amygdala or Subgenual cingulate. ‘FC’ seeds are the primary regions of interest; ‘Networks’ represent the networks from independent FC based atlases. ‘30’ represents 30+% probabaility from a histological atlas. ‘CMA’ and ‘SF’ are centromedial and superficial amygdala, respectively. ‘*’ indicates p<.05 significance; ‘+’ is p=0.080.

The sgACC region moved in the negative direction (PE=-0.073, Wald χ^2^=3.07, p=0.080) in response to targeted TMS (Figure 4). Stimulation sites with greater relative pain ratings also drove the sgACC down to a greater degree (pain PE=-0.048, Wald χ^2^=5.86, p=0.015). Dilating the mask by 1 voxel (2mm space; spherical kernel) weakened the strength of the PE (−0.036), the Wald statistic (χ^2^=1.21) and the p-value of the intercept (p=0.271), suggesting that the targeted region, and not the adjacent cortical space, was particularly affected by TMS. Testing specificity by using an independent surface based atlas for the sgACC (34), we found a similar direction in the PE (−0.040) that was weaker and was not significant (Wald χ^2^=1.69, p=0.194). Relative pain was again significantly associated with a negative average response within this ROI (PE=−0.042, Wald χ^2^=4.90, p=0.027), suggesting a broader regional effect of pain in ventromedial prefrontal cortex. It is noteworthy that both sgACC masks are in close proximity to a medial prefrontal canonical default mode network (DMN) subregion which prompted additional exploratory analyses described in a separate section below.

### Network-level responses indicated that sgACC but not amygdala networks responded to TMS

The average BLA network effect was not significant (PE=0.006, Wald χ^2^=0.29, p=0.590; Figure 4) which also was not affected by expanding the mask to z>=0.2 (PE=0.005, Wald χ^2^=0.42, p=0.518) or by using the network mask based on a gyral atlas defined amygdala as seed (z>=0.3; PE=0.005, Wald χ^2^=0.18, p=0.670). This set of findings suggests that the prefrontal-amygdala causal connection is specific to the amygdala only, rather than its distributed network. The specificity of this effect is perhaps related to direct prefrontal-amygdala links established previously in non-human primate and human studies (10-12).

For the sgACC network, there was evidence for a negative average BOLD response (PE=−0.045, Wald χ^2^=11.77, p=0.0006; Figure 4) that persisted when using the more liberal mask (PE=−0.044, Wald χ^2^=6.77, p=0.009). Using the FreeSurfer sgACC seed atlas, the network negative BOLD response was again significant and of a similar magnitude (PE=−0.045, Wald χ^2^=11.58, p=0.0006), suggesting that the sgACC network defined in at least two ways is downregulated by TMS focused on an sgACC target. Given sgACC hyperactivity in depression, this negative BOLD response is in the presumably clinically useful direction and thus lends support to the potential mechanism of neuromodulation treatment focused on this network.

The sgACC network largely overlaps with the canonical default mode network (DMN; Supplementary Figure 3) particularly in the ventromedial prefrontal cortex. We therefore conducted additional analyses aimed at addressing the specificity of the sgACC network effect with respect to DMN.

### The sgACC network effect remained when the DMN was excluded

As suspected, sgACC focused stimulation caused a deactivation in the DMN on average relative to other areas of the brain (PE=−0.028, Wald χ^2^=9.49, p=0.002; Figure 4). Masking out the overlap between the DMN from the largest (7 network) Yeo atlas (40) from the quantile images and re-fitting the GEE on the subsequent average network quantiles did not eliminate the sgACC network response to TMS (PE=−0.040, Wald χ^2^=4.04, p=0.044). Using the gyral atlas defined sgACC region network, the evoked network effect again remained significant (PE=−0.035, Wald χ^2^=5.58, p=0.018). Thus, TMS targeting the sgACC also influences the DMN but the observed significant response in the sgACC is partially independent of this effect.

### Covariates did not influence the results

Head motion, pain, and age were not significant predictors of subcortical ROI response or network response evoked by TMS in the analyses described above other than those mentioned for pain/discomfort.

### Results were robust to the choice of working correlation structure

Given the type of data, which included multiple stimulation sites per subject, we specified a working exchangeable correlation structure, which estimates a single correlation parameter for any two repeated measurements within a subject. However, as a test of sensitivity to the choice of working correlation, we repeated the GEE analyses with an independent working correlation structure, which assumes repeated measures within subjects are not correlated. Using the independent correlation structure only slightly altered parameter estimates and p-values with only two changes in statistical significance at the p<0.05 level: 1) the superficial amygdala evoked response was significant at p=.033 when using the exchangeable working correlation and was marginally above statistical significance with p=.055 when using the independent correlation model; and 2) the gyral defined sgACC pain response changed from being significant with a p=0.027 to non-significant with a p=0.067 when specifying an exchangeable and independent correlation structure, respectively. In total, the results were highly robust to the choice of correlation structure for the GEE.

### Physiological arousal models

A number of subjects were missing one or more of these physiological covariates, leading to a reduction in sample size for these tests when using a complete case analysis. However, we explored preliminarily whether these non-neuronal physiological variables were contributors to our primary results. As the amygdala in some fMRI studies covaries with autonomic arousal (41), we tested the possibility that amygdala response in our models reflected autonomic co-activation by including heart rate and respiration as additional covariates to the models with age, pain and head motion. We also added heart rate and respiration as additional covariates in the sgACC analyses to test the possibility that the cutaneous sensation of TMS drove an autonomic response which drove an sgACC response. Average heart rate and respiration derived from the pulse oximeter recorded during each TMS/fMRI run were recorded and judged to be of high quality in 33/44 runs for amygdala targets and 31/41 runs for sgACC targets. For the 30% and up BLA mask and CMA mask, neither heart rate nor respiration were significantly associated with amygdala response (BLA30 ps>0.11; CMA30 ps>0.54). For the SF 30%+ probability subregion, increased heart rate was associated with higher amygdala response (PE=0.069, Wald χ^2^=3.99, p=0.046), and the relationship remained significant with a higher SF 50% probability mask (PE=0.088, Wald χ^2^=4.10, p=0.043). The independent Freesurfer amygdala region response was not significantly associated with either physiological variable (HR p=0.90; Resp p=0.85).

In the amygdala network (BLA z>0.3), heart rate was positively associated with response to TMS (PE=0.016, Wald χ^2^=4.67, p=0.031). An even stronger relationship was found using the independent amygdala network mask (PE=0.028, Wald χ^2^=9.55, p=0.002). Again, neither network responded significantly to TMS in the original analyses.

Testing for specificity using our primary sgACC region of interest, neither of the physiological measures were significantly associated with the TMS evoked response (p>0.39) nor were they significantly associated with the independent sgACC region response (ps>0.50). Similarly, the sgACC network response was not significantly associated with either physiological variable (ps>0.31).

In summary, the amygdala network and superficial amygdala subregion responses covaried with heart rate whereas centromedial and basolateral amygdala activation were not predicted by heart rate. None of the sgACC focused results were predicted by autonomic measures.

### Exploratory voxelwise analyses

Given that the amygdala BLA was used as a target but the significant amygdala evoked effects from TMS targeting this region were most pronounced in the CMA and superficial zones, we performed a voxelwise exploratory analysis using a false discovery rate of q<0.05 within a mask that included all three subregions (overlapping) at a 30%+ threshold. The peak corrected voxel in that mask was at MNI x,y,z (−20, −6, −12), q=0.0000023 with the following histology based probabilities: 78% superficial, 68% laterobasal, and 56% centromedial. In other words, the amygdala subregions were all well represented in the peak TMS evoked amygdala response.

An average TMS evoked map from the GEE (mean + 1.5 SD) for illustration purposes for each target (sgACC and BLA) is included as Supplementary Figures 4 and 5. The thresholded average evoked TMS images demonstrate overlap with the sgACC and BLA networks defined using the independent atlas.

### Verifying quantile directional effects in contrast estimate BOLD

Though we argue in the Methods Section for the utility of converting first level GLM (TMS event relative to baseline) contrast estimates to individual quantile maps, we interpret the direction of the quantile effects (greater or less than the median (0.5)) as ‘positive’ and ‘negative’ responses, respectively, in the same way that positive or negative BOLD responses are generally interpreted. To validate this interpretation in light of the prior literature, we averaged the original contrast estimate maps across subjects and stimulation sites, extracted a mean value for individual masks of interest, and found that, indeed, there was a negative BOLD response in the sgACC network average, Mean(SD): −0.105(.053) and in the sgACC average ROI, −0.100(.027). Also consistent with our interpretation of the positive amygdala response to focused TMS, the CMA and SF regions had positive BOLD contrast weights (CMA=.034(.047); SF=.027(.069)).

Though not significant in the primary quantile analyses, the BLA network response was very slightly negative -.002(.040) as was the BLA ROI response -.008(.044). At the voxelwise amygdala response peak (Figure 5), the BOLD response was positive 0.100(0.371) as were the responses in the neighboring voxels and in the homologous contralateral amygdala (Supplementary Figure 4).

**Figure 5.**
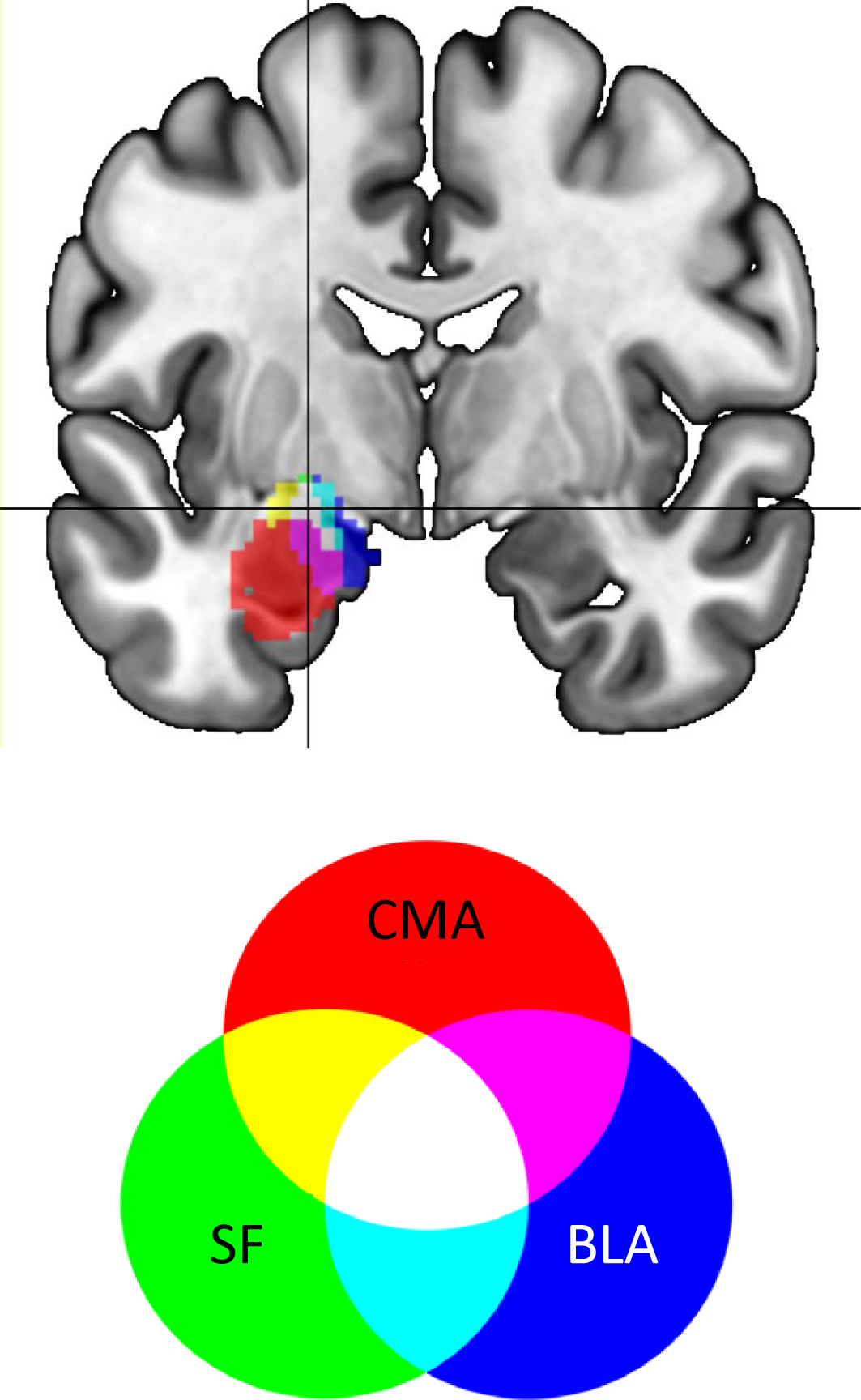
Peak amygdala response to TMS. In a combined basolateral (blue), centromedial (red) and superficial (green) amygdala volume, the peak TMS evoked brain response (FDR corrected; p<.05) at MNI xyz (−20, −6, −12) is indicated by the crosshairs and represents a coordinate with 78% SF, 68% BLA and 56% CMA probability based on an histology generated atlas.

## Discussion

In this investigation, we provide the first evidence that individually targeted TMS can indirectly and noninvasively engage subcortical areas and their distributed network representations. TMS treatments are delivered to single brain areas but their clinical effects are not likely confined to the stimulation site. Focusing on pairs of brain areas as stimulation sites and downstream targets allows proof-of-principle support for targeted approaches to brain stimulation, as we establish here using TMS to target the subgenual anterior cingulate cortex (sgACC) and amygdala via prefrontal stimulation sites. A narrow set of circuits may be critical for effecting clinical changes and the neuroimaging literature may be an appropriate guide suggesting the existence and location of these circuits. A potentially more behaviorally meaningful approach is discovering the degree to which a particular network node might causally influence a brain network. We have established this approach for targeting canonical resting state brain networks (1) that we build upon here demonstrating that the sgACC as well as the distributed functional network with which it is connected respond to individualized sgACC targeted TMS. The sgACC response was effective in moving the target region but did not move an adjacent region generated using a surface parcellation atlas. Though both regions had zones of negative BOLD response, only the sgACC target mask had a homogenous negative response whereas the surface based region showed mixed positive and negative peaks. The amygdala response peak was in a zone where several subregions of the amygdala complex overlap as determined by a human histological atlas (29). The amygdala responded to targeted TMS though the distributed amygdala network was not effectively engaged. This pattern wherein the surface node and downstream target only were engaged suggests a tight prefrontal-amygdala coupling separate from each node’s distributed brain network profile.

In this investigation of resting fMRI guided targeting, we determined that this approach was effective given that we causally influenced both targets by applying TMS to prefrontal sites determined using resting fMRI. The precise targeting location varied by individual as did each subject’s TMS evoked brain map that was ranked voxelwise by quantile before being aggregated for sgACC or amygdala target analyses. In the behavioral domain, it has been demonstrated that individual functional localization is better than anatomical localization which is better than a group functional template which is better than a scalp EEG coordinate in terms of effect size influences on behavior (42). Our results are consistent with these preliminary findings and support recent cognitive neuroscience work that observes individually specific functional network mapping (2, 3).

TMS targeting was done using mostly positively correlated prefrontal sites with subcortical targets of interest. Nevertheless, the stimulation sites differed in the direction of induced BOLD response with the amygdala target generating a positive BOLD response and the sgACC generating a negative BOLD response. The hemodynamic response to TMS likely depends on a number of factors including the degree to which inhibitory interneurons are activated (43), whether the induced field is strong enough to activate synapses in addition to axons (44) and also the brain state at the time of stimulation (45). It has been shown, for example, that TMS to one area of the visual system (frontal eye field) results in a BOLD decrease in the central visual field with a concomitant increase in peripheral field brain representations (46). The strength of single TMS pulses can be influenced by the brain state such as when the effects of visual adaptation are counteracted (47) or when the brain is engaged in processing sensory information (45) as well as during mental states such as motor imagery (48, 49), anxiety (49), and so on. Manipulating and/or measuring brain state during TMS will add additional understanding to the conditions favoring specific directional changes in BOLD response.

Seeding the sgACC in previous studies of resting fMRI has generally found the distributed default mode network (DMN) map (50). Individual differences in sgACC connectivity were also reproducible across scan sessions. Using our analysis pipeline, we found substantial overlaps between the sgACC network and the canonical DMN (40) as well as a DMN response to sgACC focused TMS. When removing the sgACC network voxels that overlapped with the largest DMN mask (Yeo 7 Network), we continued to find an sgACC network response, suggesting the induced brain response to sgACC targeted TMS includes a brain response not captured by the DMN.

Neither the sgACC, the sgACC network nor the centromedial amygdala subregion responses to TMS were predicted by variations in heart rate or respiration recorded during TMS/fMRI runs. One exception was the heart rate relationship with the superficial amygdala response that was unique to that subregion. We also found that, though the amygdala network was generally not responsive to TMS, there was a significant relationship between amygdala network response and heart rate, suggesting an interdependence between the amygdala network and cardiovascular response to TMS.

Limitations in the present study included a relatively small number of participants (though a large number of stimulation sites across participants). Also, though the sgACC network was partially independent of DMN response to TMS, the sgACC ROI and sgACC network at least partially covaried with the DMN response and so DMN response to TMS could contribute mechanistically to, for example, repetitive TMS (rTMS) treatment in depression shown to alter sgACC-DMN connectivity (27). We here shed light on a targeting approach for administering TMS to engage sgACC and amygdala. However, we do not yet establish the degree to which these circuits are modifiable using rTMS or other interventions. Finally, the selection of a target for an individual participant among a variety of ‘hot spots’ indicated by the resting fMRI map will require additional focused study especially in neuromodulatory investigations.

## Conclusion

We here establish that resting fMRI guided non-invasive brain stimulation is effective in causally and specifically influencing subcortical targets such as the subgenual anterior cingulate cortex and the amygdala. Adding to enthusiasm for ‘individualized’ brain network representations, we demonstrate the value of individualizing TMS target locations for each participant and individualizing brain responses through analysis of quantile values taken from 1^st^ level GLM contrasts in TMS-evoked fMRI data. We further contend that TMS/fMRI is a powerful technique for probing causal circuit pathways that can uncover mechanistic details especially in determining how neuromodulation changes brain, behavior, and symptoms. Future research in our lab and others will establish the relevance of this type of circuit mapping approach to individual differences in response to neuromodulation targeting specific brain networks. The ultimate goal of TMS mapping and neuromodulation studies in healthy and patient populations is to optimize neuromodulation in order to move the brain from a less to a more optimal state (51). The field of interleaved TMS/fMRI is still in its infancy and approaches to use these causal maps to investigate neuropsychiatric abnormalities is worthy of focused study in its own right (52-54).

## Acknowledgements

We thank Dr. Yvette Sheline for contributions to this project including equipment, scan time and personnel. We also thank Yordan Todorov of Magventure, Inc. for TMS device support and contributing customized equipment for hardware control.

## Supplementary material

Supplementary material is available online.

## Supplementary Material | Oathes *et al*

**Supplementary Figure 1.**
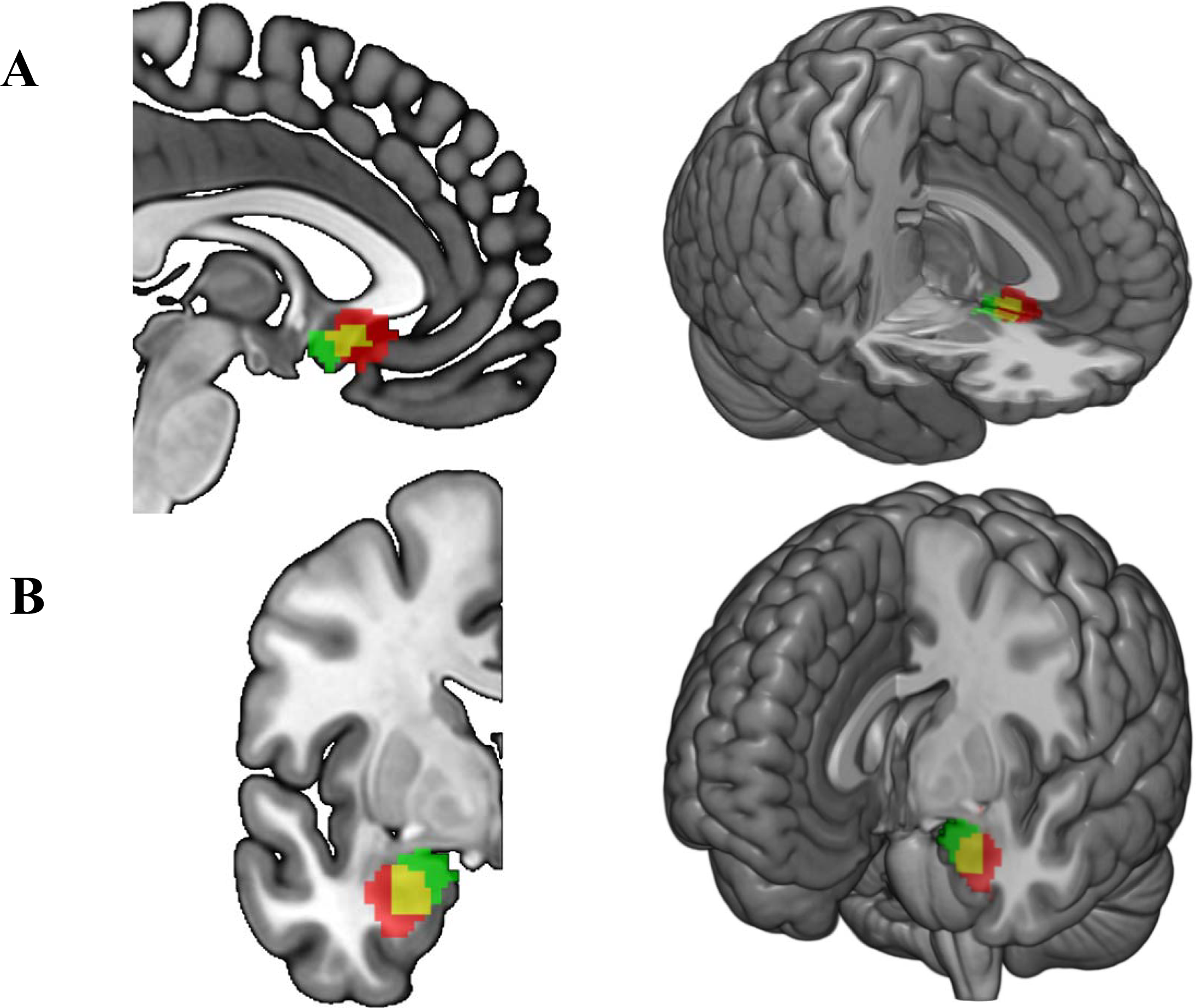
FreeSurfer vs. primary regions of interest / functional connectivity seeds. (**A**) FreeSurfer (Green) overlapping (Yellow) with primary (Red) subgenual anterior cingulate regions of interest. (**B**) FreeSurfer (Green) overlapping (Yellow) with primary (Red) amygdala regions of interest.

**Supplementary Figure 2.**
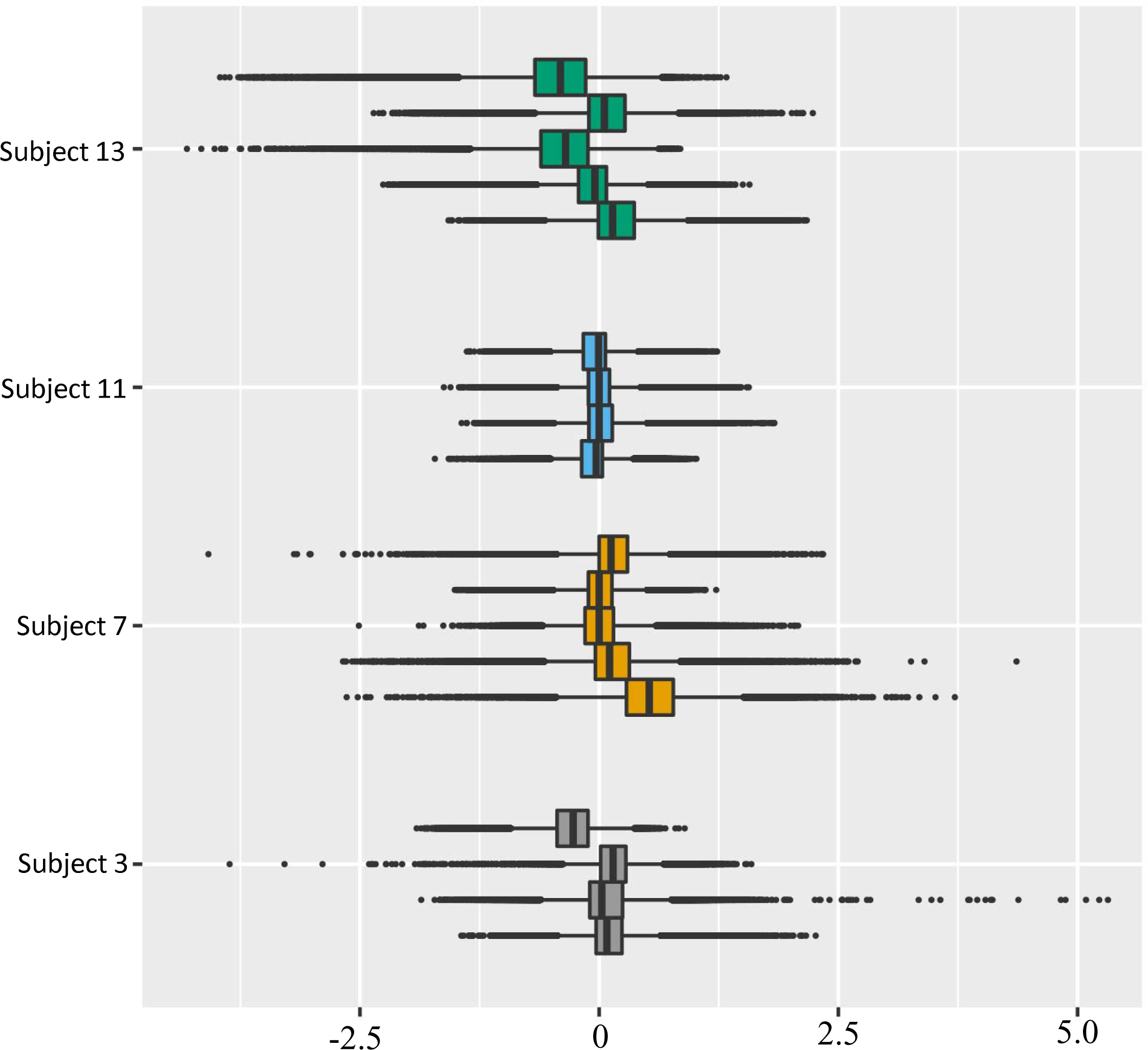
Variability in TMS evoked brain responses. To illustrate the motivation for converting fMRI BOLD contrast estimates to quantiles and for using generalized estimating equations accounting for cross-subject variability, TMS evoked contrast estimates for each site within a subject (color coded) are shown according to each voxel in the brain shown as a single point (black dots). Solid lines in the bars represent the median and the box hinges are the 25^th^ and 75^th^ percentiles. Whiskers are at the min/max but no further than 1.5* IQR (interquartile range).

**Supplementary Figure 3.**
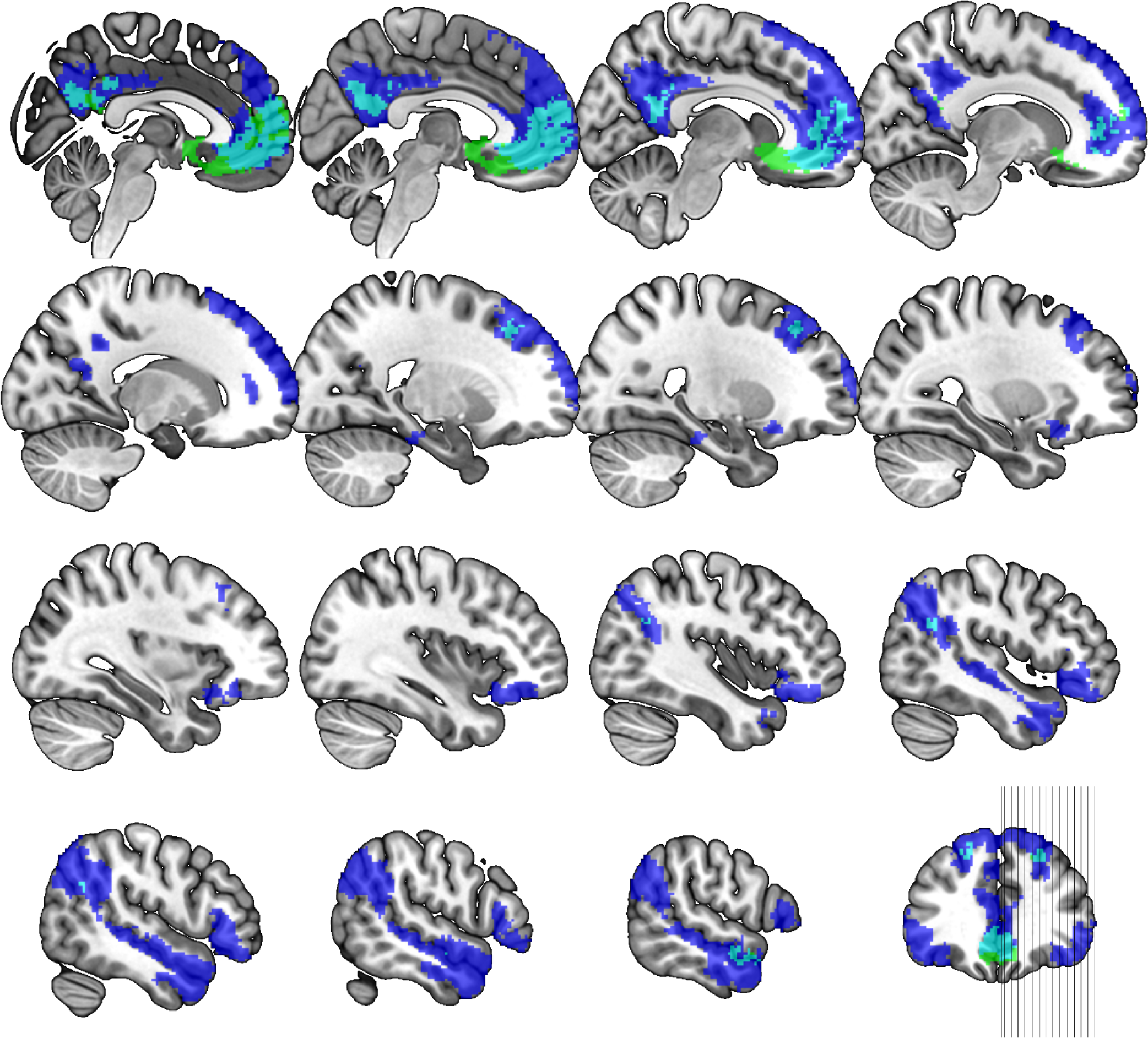
Subgenual cingulate vs. DMN network. Using an independent data set, the network mask shown was created seeding the primary (green) defined sgACC and calculating functional connectivity (correlation) with z≧0.3 in 96+/127 subjects, effect size >6.0 and cluster ≧ 2mm3 in 2mm MNI standard space (excluding seed). The Yeo 7 network DMN mask is shown in dark blue and the overlap between the sgACC network and DMN are in light blue. Slice wise views are represented starting at MNI x=2 and moving out in 2mm steps with an extra step between rows until the final x=58 image.

**Supplementary Figure 4.**
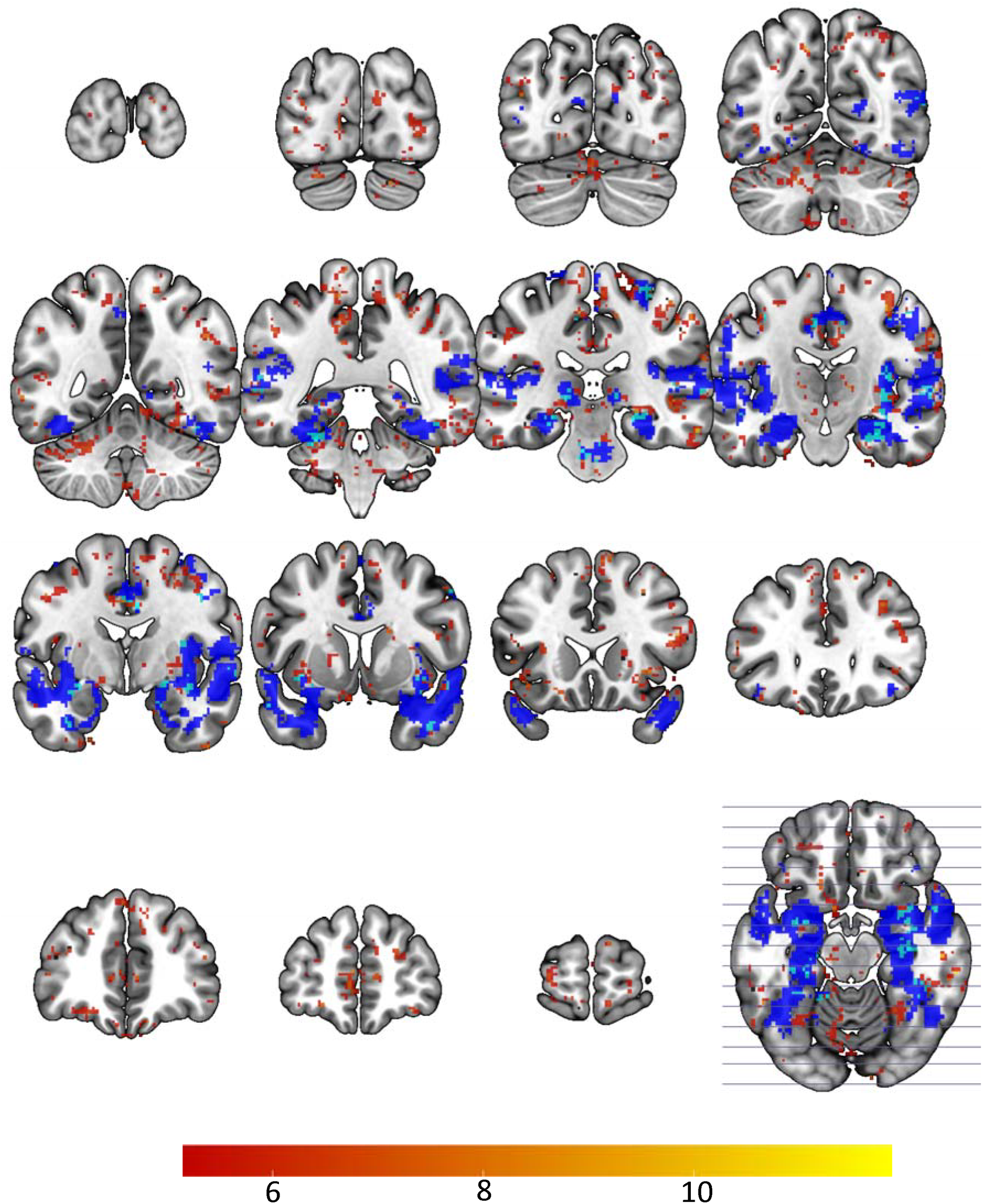
Average Wald Chi-Sq of the BLA TMS evoked map. Basolateral amygdala network mask (dark blue) with TMS evoked quantile mask (mean + 1 standard deviation; red) with their overlap in light blue. Slice wise views are represented starting at MNI y=−98 and stepping from right to left from the top left at y=−86, −64; −52, −40, −28, −16 (bottom right at y= −16).

**Supplementary Figure 5.**
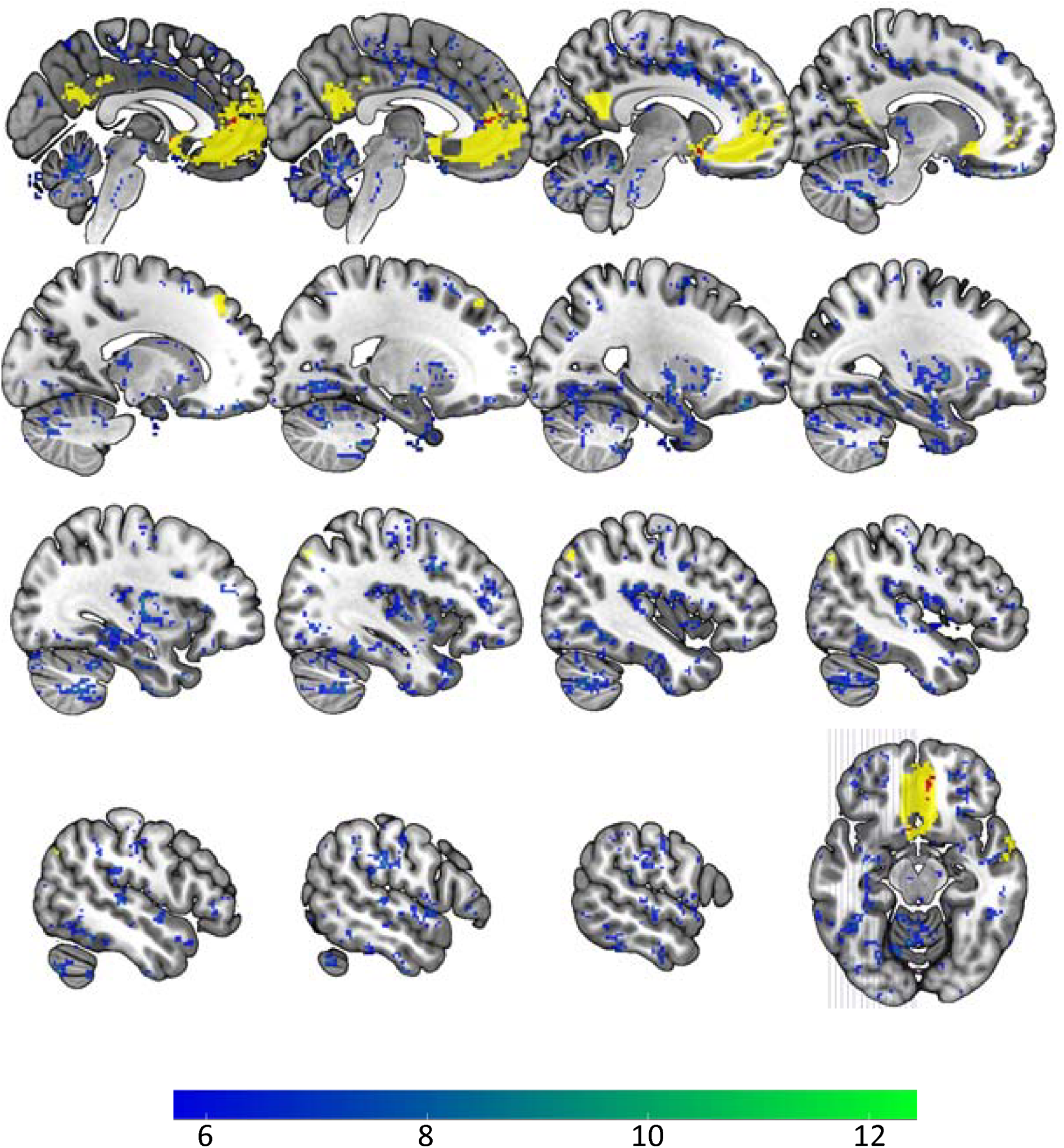
Average Wald Chi-Sq of the sgACC TMS evoked map. Subgenual anterior cingulate network mask (yellow) with TMS evoked quantile mask (mean + 1 standard deviation; blue/green) with their overlap in red. Slice wise views are represented starting at MNI x=-2 and stepping from right to left from the top left at x=−4, −8; −12; −16, −21, −25, −29; −33, −37, −42, −46; −50, −54, −58 (bottom right at x= −14).

